# Scarless conditional sgRNAs via endogenous mascRNA processing enable rapid and temporally controlled genome editing

**DOI:** 10.64898/2026.08.20.745814

**Authors:** Curtis Hart, Lovely Paul Solomon Devakumar, Khalid Saeed, Adam Spruce, Chara Mastrokalou, Sebastian Lukasiak, Douglas Ross-Thriepland, David Walter, Nikhil Gupta

**Affiliations:** Cancer Research Horizons, Cancer Research UK, Cambridge, United Kingdom; Discovery Sciences, BioPharmaceuticals R&D, AstraZeneca, Cambridge, United Kingdom

**Author notes:** These authors contributed equally to this work (Curtis Hart, Lovely Paul Solomon Devakumar). King Abdullah International Medical Research Centre, King Saud Bin Abdulaziz University for Health Sciences, Riyadh, Saudi Arabia (Khalid Saeed). Cambridge Stem Cell Institute, University of Cambridge, Cambridge, United Kingdom (Adam Spruce). Department of Molecular Biology and Genetics, Faculty of Science, Bilkent University, Ankara, Türkiye (Nikhil Gupta).

## Abstract

Precise temporal control of gene editing is essential for studying dynamic biological processes, interrogating essential gene function, and improving the interpretability of pooled perturbation screens. Cre-dependent single guide RNA (sgRNA) switches provide temporal regulation by coupling guide activation to site-specific recombination, but existing designs retain a loxP-derived 5′ sequence (scar) on the mature sgRNA that can impair guide function. We developed a scarless conditional sgRNA platform that combines Cre-loxP recombination with endogenous RNA processing to restore the native sgRNA architecture following induction. A MALAT1-associated small cytoplasmic RNA (mascRNA) module was positioned upstream of the guide sequence such that, after Cre-mediated recombination, cellular RNase P and RNase Z remove the residual loxP-derived overhang, generating a mature sgRNA with an authentic 5′ terminus. Using guides targeting endogenous cell-surface marker genes, the scarless design maintained stringent OFF-state control while improving ON-state editing performance compared with a conventional Cre-activated sgRNA switch, resulting in faster editing kinetics, greater perturbation penetrance, and more consistent editing efficiency. This modular strategy provides a simple approach for conditional CRISPR genome editing that preserves guide integrity and should be readily adaptable to time-resolved functional genomics and pooled screening applications.

## Introduction

The CRISPR-Cas9 system has transformed genome engineering by enabling precise, programmable DNA cleavage across diverse cellular contexts (Jinek et al., 2012; Cong et al., 2013). Beyond reverse genetics, CRISPR-based perturbation underpins genome-scale functional screens that identify gene essentiality, therapeutic vulnerabilities, and regulatory mechanisms in human disease (Shalem et al., 2014; Wang et al., 2014; Doench, 2018). However, conventional CRISPR systems lack the precise temporal control of editing required to study dynamic biological processes or essential genes, where constitutive disruption leads to rapid negative selection before gene function can be interrogated at defined biological time points (Hart et al., 2015; Doench, 2018). Temporal regulation is therefore essential for investigating processes such as cell-fate transitions, stress responses, and disease progression, in which gene perturbation must be synchronized with specific biological events (Zhu et al., 2016; Norman et al., 2019).

Two broad strategies have been developed to control CRISPR activity: inducible Cas9 expression and inducible sgRNA activation. The most widely used Cas9 systems employ doxycycline-responsive Tet-On promoters or small-molecule-regulated destabilization domains (González et al., 2014; Dow et al., 2015; Zetsche et al., 2015; Senturk et al., 2017). While effective, these approaches frequently exhibit basal Cas9 activity in the uninduced state, resulting in leaky editing that complicates studies of essential genes and transient cellular phenotypes (Srinivasan et al., 2026). An alternative strategy regulates the sgRNA rather than Cas9 (Aubrey et al., 2015). Cre-loxP-dependent sgRNA switches exploit the high efficiency and versatility of Cre recombination by placing a loxP-flanked transcriptional stop cassette (polyT sequence) upstream of the guide sequence (Sternberg and Hamilton, 1981; Chylinski et al., 2019). Cre-mediated excision activates sgRNA expression and is readily achieved through transient Cre mRNA electroporation, recombinant Cre protein delivery, or inducible Cre transgenes, making these systems compatible with existing Cas9-expressing models (Indra et al., 1999; Jullien et al., 2003; Van den Plas et al., 2003).

Despite these advantages, current Cre-activated sgRNA designs retain a 34-nucleotide loxP sequence at the 5′ end of the mature guide following recombination (Chylinski et al., 2019). Although the functional consequences of this residual sequence have not been directly examined, biochemical studies have shown that 5′ guide extensions and altered secondary structure can impair Cas9 ribonucleoprotein assembly, alter R-loop formation, and reduce DNA cleavage efficiency in a sequence-dependent manner (Briner et al., 2014; Mullally et al., 2020). In addition, guide activity is inherently influenced by spacer sequence and mismatch tolerance (Hsu et al., 2013). Together, these observations suggest that the residual loxP sequence may delay editing kinetics, reduce perturbation penetrance, and contribute to guide-to-guide variability, representing an underappreciated limitation of conditional CRISPR systems.

RNA processing offers a potential solution by restoring the native sgRNA architecture after transcription. Endogenous RNase P and RNase Z have been widely exploited in tRNA based multiplex CRISPR systems to release mature sgRNAs with defined 5′ termini from polycistronic transcripts (Xie et al., 2015; Port and Bullock, 2016; Dong et al., 2017). However, canonical tRNAs contain internal RNA polymerase III promoter elements that drive constitutive transcription, rendering them unsuitable for conditional guide systems because they compromise OFF-state control (Knapp et al., 2019).

MALAT1-associated small cytoplasmic RNA (mascRNA) provides an attractive alternative. mascRNA is a naturally occurring 58-nucleotide tRNA-like RNA generated from the 3′ end of the long non-coding RNA MALAT1 through sequential processing by RNase P and RNase Z (Wilusz et al., 2008; Arun et al., 2020; Skeparnias et al., 2024). Unlike canonical tRNAs, mascRNA lacks intrinsic RNA polymerase III promoter activity while retaining efficient recognition by the endogenous tRNA-processing machinery, making it well suited as an RNA processing module within inducible sgRNA architectures (Knapp et al., 2019).

Here, we incorporate a mascRNA processing module immediately upstream of the sgRNA spacer within a Cre-loxP conditional guide cassette. Following Cre-mediated recombination, endogenous RNase P cleaves upstream of the tRNA-like fold to remove the residual loxP-derived 5’ overhang, and RNase Z cleaves at the mascRNA 3’ end, restoring the native sgRNA terminus. We show that this scarless design maintains stringent OFF-state control while improving the efficiency, kinetics, and consistency of genome editing relative to a conventional Cre-activated sgRNA switch, providing a modular platform for temporally controlled genome editing and functional genomics.

## Methods

### Cell lines and culture conditions

HT-29 and RKO colorectal adenocarcinoma cells constitutively expressing Streptococcus pyogenes Cas9 (SpCas9) were used throughout this study (HT-29_Cas9 and RKO_Cas9). Stable Cas9 expression was generated by lentiviral transduction of pKLV2-EF1a-Cas9Bsd-W (Addgene #68343) in the presence of 8 µg/ml polybrene (Merck), followed by selection with 10 µg/ml blasticidin for 4 days. Cas9 expression and nuclease activity were validated by assessing fluorescent protein loss following transduction with pKLV2-U6gRNA5(gGFP)-PGKBFP2AGFP-W (Addgene #67980). HT-29_Cas9 cells were maintained in RPMI-1640 medium, while RKO_Cas9 cells were maintained in MEM (Eagle’s). All media were supplemented with 10% (v/v) fetal bovine serum (Gibco, A5256701), 1X GlutaMAX (Gibco, 35050061), and 1X penicillin– streptomycin (Gibco, 15140122). Cells were cultured at 37°C in a humidified incubator containing 5% CO_2_.

### Construct design and cloning

Guide RNA sequences targeting CD151, B2M, and AAVS1 were designed using the VBC guide-design tool (Vienna BioCenter; vbc-score.org) (Michlits et al., 2020). All guide constructs were assembled into an in-house lentiviral backbone driven by the human U6 RNA polymerase III promoter. Constitutive guide constructs consisted of: (i) U6-guide, containing an unmodified sgRNA sequence; (ii) U6-loxP-guide, containing a single loxP sequence immediately upstream of the guide; and (iii) U6-loxP-mascRNA-guide, containing a single loxP sequence followed by a mascRNA processing module and guide sequence. Inducible guide constructs consisted of: (iv) U6-loxP-polyT-loxP-guide, containing a loxP-flanked transcriptional termination cassette upstream of the guide; and (v) U6-loxP-polyT-loxP-mascRNA-guide, containing the mascRNA processing module between the residual loxP site and guide sequence following recombination. A tRNA-containing conditional construct was generated by replacing the mascRNA module with a 71-nucleotide minimal fly-tRNA sequence to assess the impact of endogenous Pol III promoter activity on OFF-state control. The mascRNA sequence was derived from the human MALAT1 locus and incorporated the 58-nucleotide mascRNA processing element together with the surrounding sequences required for recognition and cleavage by endogenous RNase P and RNase Z. Transcriptional termination sequences consisted of consecutive thymidine residues encoded within the DNA template. All constructs were synthesised by Genewiz and verified by Sanger sequencing prior to lentiviral production. Guide sequences used in this study are provided in Supplementary File 1.

### Lentiviral production and transduction

HEK293T cells were seeded at 1.1 × 10^6^ cells per well in six-well plates approximately 24 h before transfection and cultured until reaching 80–90% confluency. Cells were co-transfected with lentiviral transfer plasmids (1.65 µg), psPAX2 packaging plasmid (1.37 µg), and pMD2.G envelope plasmid (0.55 µg) using Lipofectamine 3000 (5.21 µl; Thermo Fisher Scientific, L3000001). Medium was replaced 24 h after transfection, and viral supernatants were collected 72 h after transfection, clarified using a 0.45 µm syringe filter, and stored at −70°C until use. Target cells were transduced with filtered viral supernatant in the presence of 8 µg/ml polybrene at a multiplicity of infection (MOI) of approximately 0.3 to favour single-copy integration events. Four days after transduction, cells were selected with puromycin (3 µg/ml for HT-29_Cas9 and 0.6 µg/ml for RKO_Cas9) for 4 days.

### Cre recombinase delivery

Transient Cre-mediated recombination was achieved by electroporation of synthetic Cre recombinase mRNA (Cre-mRNA; Trilink Biotechnologies, L-7211). A total of 5 × 10^6^ cells were electroporated with 1 µg Cre mRNA using a MaxCyte electroporation system with an OC-100x2 processing assembly. Mock-electroporated cells, which underwent electroporation without Cre mRNA, were used as uninduced controls. Following electroporation, cells were recovered in antibiotic-free medium for 24 h before returning to standard culture conditions.

### Flow cytometric analysis

Surface expression of CD151 and B2M was quantified by flow cytometry at days 0, 4, 7, 11, and 14 following transduction (constitutive constructs) or Cre mRNA electroporation (inducible constructs). At each timepoint, cells were detached using Accutase, fixed in 4% (w/v) paraformaldehyde for 20 min at room temperature, washed with phosphate-buffered saline containing 2% fetal bovine serum, and stored at 4°C. To reduce inter-day variation, all samples within each biological replicate were subsequently stained together in a single batch with fluorophore-conjugated antibodies for 30 min at 4°C. CD151 depletion was measured using PE-conjugated anti-human CD151 antibody (BioLegend, 350408). B2M depletion was assessed indirectly using FITC-conjugated anti-human HLA-A,B,C antibody (clone W6/32; BioLegend, 311404), which recognises a conformational epitope dependent on B2M association. Antibodies were used at 1 µl per 100 µl staining reaction. Samples were acquired using a MACSQuant flow cytometer (Miltenyi Biotec) and analysed using FlowJo version 10.10.0. All experiments were performed using biological triplicates unless otherwise indicated.

### Assessment of tRNA-based processing constructs

To evaluate the suitability of tRNA as an RNA processing module in a conditional sgRNA system, HT-29_Cas9 cells were transduced with either the standard inducible U6-loxP-polyT-loxP-guide construct or a U6-loxP-polyT-loxP-tRNA-guide construct targeting CD151. Constructs targeting the AAVS1 safe-harbour locus were included as controls for guide-independent effects. Cells were electroporated with or without Cre recombinase mRNA, and CD151 surface expression was quantified by flow cytometry 7 days after electroporation.

### Assessment of mascRNA-based sgRNA constructs

To evaluate the effect of mascRNA-mediated processing on genome editing, HT-29_Cas9 and RKO_Cas9 cells were transduced with constitutive (U6-guide, U6-loxP-guide, or U6-loxP-mascRNA-guide) or inducible (U6-loxP-polyT-loxP-guide or U6-loxP-polyT-loxP-mascRNA-guide) sgRNA constructs targeting CD151 or B2M. Corresponding AAVS1-targeting constructs were included as controls for guide-independent effects. For constitutive guide experiments, the day of lentiviral transduction was defined as day 0. For inducible guide experiments, cells were electroporated with or without Cre recombinase mRNA, and the day of electroporation was defined as day 0. Surface expression of CD151 and B2M was quantified by flow cytometry at days 0, 4, 7, 11, and 14 to assess genome editing kinetics.

### Statistical analysis

All experiments were performed using biological triplicates unless otherwise stated, where a biological replicate is defined as an independent experiment on newly thawed cell cultures from lentiviral transduction to analysis. Data are presented as mean ± standard deviation (SD).

Two statistical approaches were used, reflecting the different experimental designs. Constitutive guide constructs, which were transduced from the same cell pool on Day0, were compared at each timepoint using a two-tailed ratio paired t-test. Inducible guide constructs, which were generated from independently transduced pools prior to Day0, were compared using a two-tailed unpaired t-test with Welch’s correction. A P-value <0.05 was considered statistically significant.

### Quantification of Cre-loxP recombination efficiency

Genomic DNA was isolated from HT-29 cells at days 0, 4, 7, 11, and 14 following transient Cre mRNA electroporation using the DNeasy Blood and Tissue Kit (Qiagen, 69504) according to the manufacturer’s instructions. The loxP-sgRNA cassette was PCR-amplified and analysed by paired-end sequencing. Thirty samples were analysed in total, representing inducible sgRNAs targeting AAVS1, B2M, and CD151, each generated with or without the mascRNA processing module across five time points. Sequencing data were analysed using CRISPResso2 (Clement et al., 2019) in batch mode. For each construct, loxP sites non-recombined and recombined amplicon sequences were provided using the reference and expected recombined amplicon settings, respectively. The recombined amplicon represented Cre-mediated excision of the polyT-loxP cassette, resulting in a single loxP sequence positioned upstream of the sgRNA architecture. Separate amplicon definitions were generated for constructs containing or lacking mascRNA. Reads failing base quality or alignment quality thresholds were excluded prior to analysis. Remaining reads were classified as non-recombined or recombined loxP sites based on alignment to the corresponding reference amplicons. The proportion of reads assigned to the recombined amplicon was used as a measure of Cre-loxP recombination efficiency. Recombination frequencies were visualised as stacked bar plots showing the proportion of non-recombined and recombined loxP sites across time points for each sgRNA construct. Sequencing data are available through BioStudies accession number E-MTAB-17181. Primer sequences used in this study are listed in Supplementary File 1.

## Results

### mascRNA-mediated processing restores native guide RNA architecture following Cre recombination

Conventional Cre-dependent sgRNA activation systems excise a loxP-flanked transcriptional termination cassette to position the U6 promoter upstream of the guide sequence (Chylinski et al., 2019). Following recombination, a residual 34-nucleotide loxP sequence remains at the 5′ end of the sgRNA, immediately adjacent to the spacer region required for efficient Cas9 ribonucleoprotein assembly and target recognition (Mullally et al., 2020). We hypothesized that inserting an RNA processing module between the loxP cassette and guide sequence would redirect this overhang into the endogenous tRNA maturation pathway, such that RNase P and RNase Z would restore the native sgRNA architecture after Cre recombination (Figure 1A).

**Figure 1.**
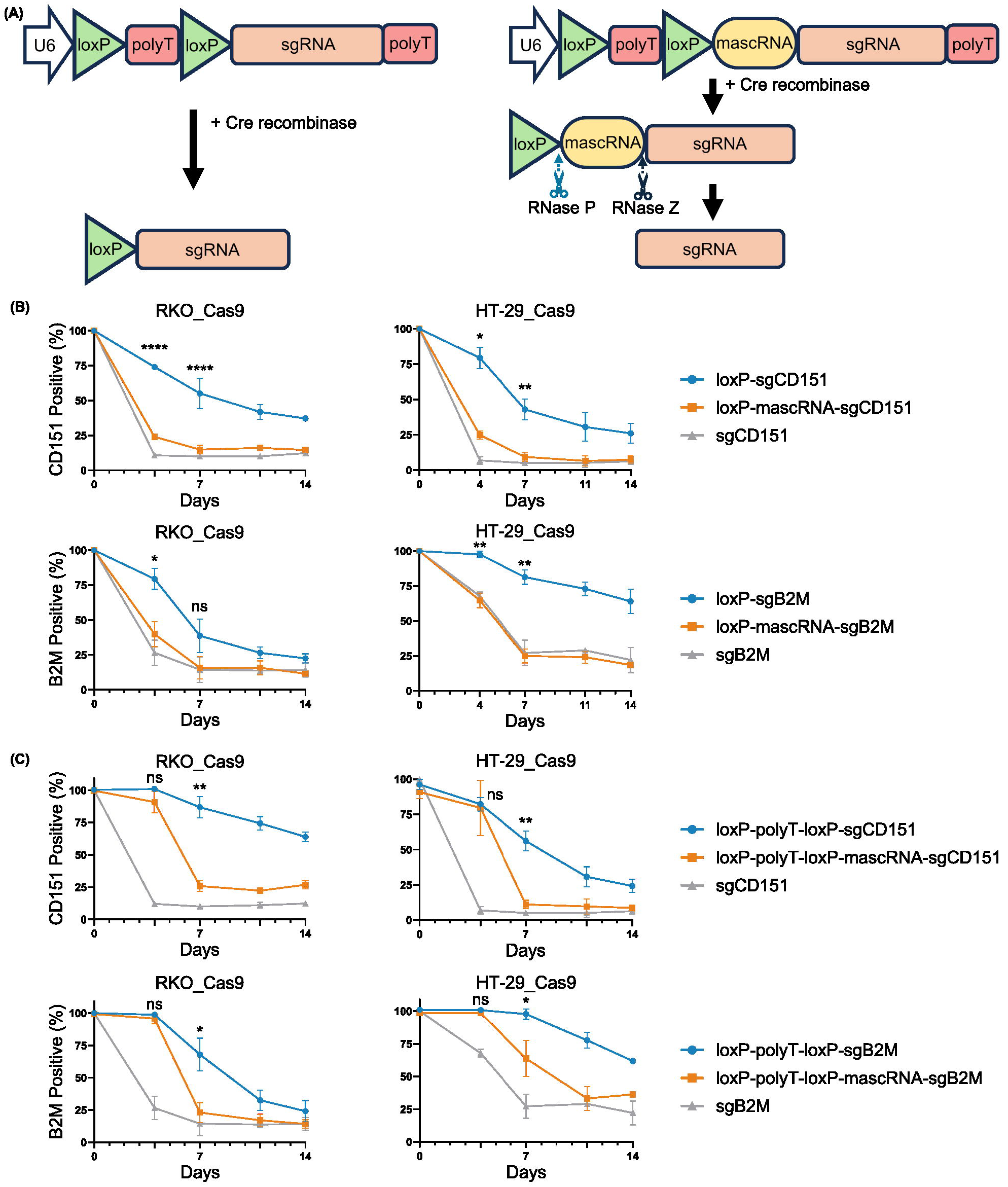
mascRNA-mediated processing restores guide architecture and improves editing kinetics in constitutive and inducible conditional systems. (A) Schematic diagram of conditional sgRNA construct architectures. Canonical Cre-inducible loxP-polyT-loxP guide; and mascRNA-incorporating inducible guide. Arrows indicate RNase P and RNase Z cleavage sites on the mascRNA element. Post-recombination processing is shown to yield a mature sgRNA with an authentic 5′ terminus. (B) Flow cytometric quantification of CD151 and B2M surface expression in HT-29 and RKO Cas9 expressing cells transduced with constitutive guide constructs. Cells were assessed at days 0, 4, 7, 11, and 14 post-lentiviral transduction. The U6-loxP-guide construct displays delayed marker loss relative to the U6-guide control, while the U6-loxP-mascRNA-guide construct restores kinetics equivalent to the unmodified control. (C) Flow cytometric quantification of CD151 and B2M surface expression in HT-29 and RKO Cas9 expressing cells transduced with the indicated respective constructs and electroporation of Cre mRNA. Time points are days 0, 4, 7, 11, and 14 post-Cre mRNA electroporation. The mascRNA-containing inducible construct achieves near-complete surface marker loss by day 7, whereas the standard conditional construct reaches only partial depletion by day 14. Data are shown as mean ± SD of three independent biological replicates. Statistical comparisons were performed using two-tailed ratio-paired t-tests between U6-loxP-guide and U6-loxP-mascRNA-guide constructs (B), and between U6-loxP-polyT-loxP-guide and U6-loxP-polyT-loxP-mascRNA-guide constructs using a two-tailed unpaired t-test with Welch’s correction (C). A P-value <0.05 was considered statistically significant. ns, not significant; *, P <0.05; **, P <0.01; ***, P <0.001.

Although tRNA sequences have been widely used to process multiplexed sgRNAs (Xie et al., 2015; Port and Bullock, 2016; Dong et al., 2017), their intrinsic RNA polymerase III promoter activity makes them unsuitable for conditional systems (Knapp et al., 2019). Consistent with this, incorporation of a tRNA sequence into a loxP-flanked guide cassette resulted in Cre-independent sgRNA activity, demonstrating loss of OFF-state control (Figure S1A).

We therefore incorporated a mascRNA processing module upstream of the guide sequence (Figure 1A). To assess the effect of the residual loxP sequence, we generated constitutive guide constructs expressing either a standard sgRNA (U6-guide), an sgRNA containing a 5′ loxP overhang (U6-loxP-guide), or the same construct incorporating mascRNA (U6-loxP-mascRNA-guide). We also generated corresponding inducible Cre-dependent guide cassettes with or without mascRNA (Figure S1B). To compare editing performance, constitutive constructs targeting the endogenous cell-surface markers CD151 and B2M were transduced into Cas9-expressing HT-29 and RKO cells. Editing kinetics were quantified by flow cytometric analysis of surface protein loss at days 0, 4, 7, 11, and 14 after transduction (Figure 1B).

The unmodified U6-guide produced rapid, near-complete loss of both CD151 and B2M by day 7 in both cell lines. In contrast, the U6-loxP-guide exhibited substantially delayed editing, with maximal depletion observed only by day 14 and remaining consistently lower than the unmodified guide (Figure 1B). These findings indicate that the residual loxP-derived 5′ sequence reduces editing efficiency, consistent with previous reports showing that 5′ guide extensions impair Cas9 activity (Hsu et al., 2013; Mullally et al., 2020).

Insertion of the mascRNA module restored editing kinetics to levels comparable with the unmodified guide. Both CD151 and B2M were efficiently depleted by day 7 in both cell lines, markedly improving editing relative to the loxP-overhang construct (Figure 1B). All datasets were normalised to AAVS1-targeting controls to account for guide-independent effects.

We next evaluated the inducible guide architectures following transient Cre mRNA electroporation. The conventional loxP-polyT-loxP-guide produced gradual and incomplete depletion of CD151 and B2M, whereas the mascRNA-containing construct achieved near-maximal editing by day 7, closely matching the kinetics of constitutive guide expression (Figure 1C). A modest delay relative to constitutive guides likely reflects the additional time required for Cre-mediated recombination and RNA processing before mature sgRNA generation. Together, these results indicate that mascRNA-mediated processing restores efficient guide function following Cre recombination.

### mascRNA-based inducible sgRNA systems preserve stringent OFF-state control

To determine whether mascRNA affected OFF-state fidelity, HT-29 Cas9 cells were transduced with conventional or mascRNA-containing inducible sgRNA constructs targeting CD151. AAVS1-targeting guides served as controls, and CD151 surface expression was monitored by flow cytometry in the presence or absence of transient Cre mRNA delivery (Figure 2A).

**Figure 2.**
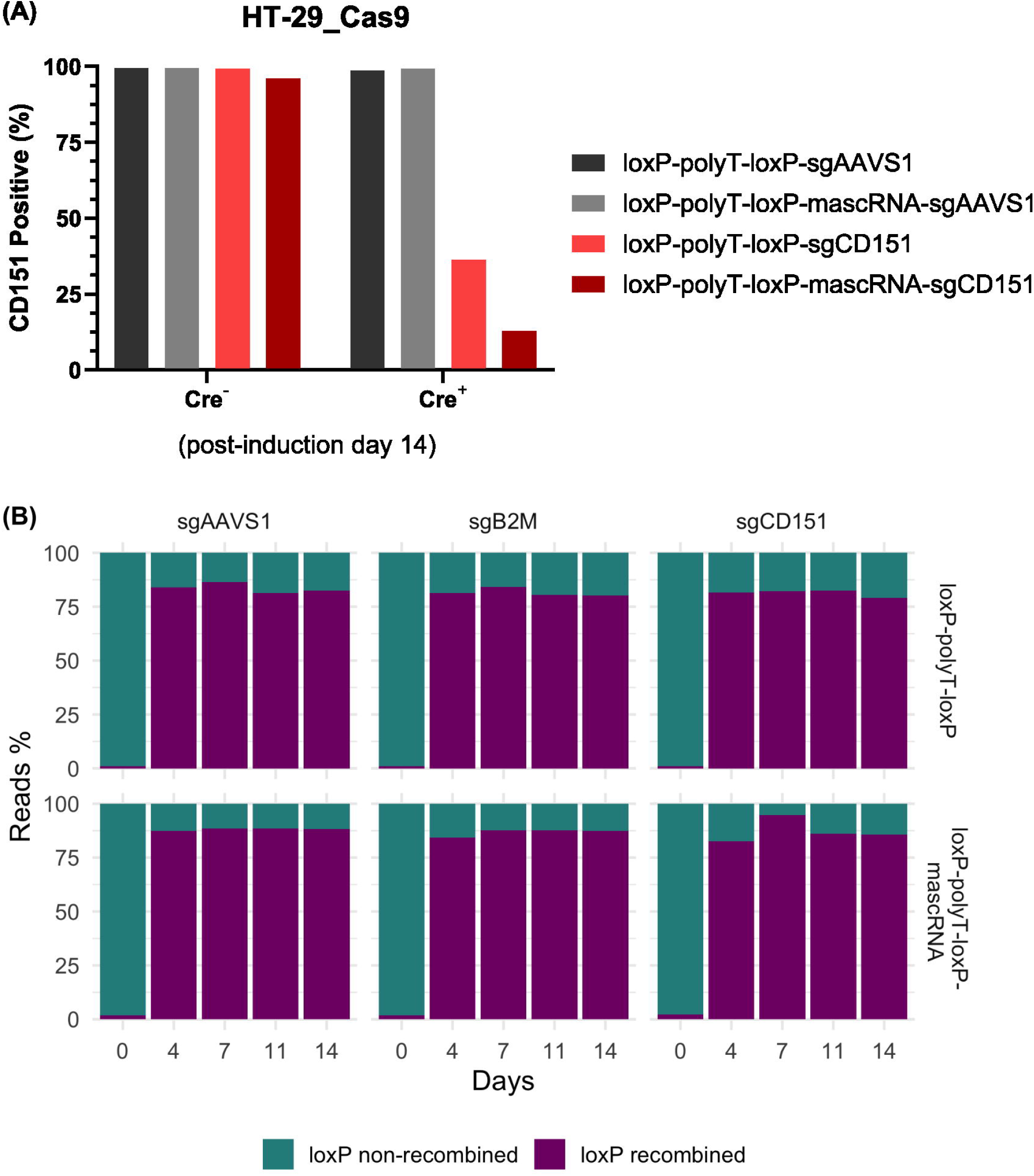
The mascRNA inducible system is non-leaky prior to Cre recombination and achieves efficient loxP-loxP recombination upon induction. (A) Leakiness assessment. HT-29 cells constitutively expressing Cas9 were transduced with inducible constructs targeting CD151 or AAVS1 (safe harbour negative control). Surface CD151 expression was monitored by flow cytometry post electroporation with and without Cre-recombinase mRNA on day14. (B) Recombination efficiency assessment. Following transient Cre mRNA electroporation, genomic DNA from HT-29 cells was harvested at days 0, 4, 7, 11, and 14 with and without mascRNA sequence, and guide cassette regions were amplified, sequenced and analysed using CRISPResso2.

In the absence of Cre, neither inducible construct produced detectable loss of CD151 expression, indicating stringent OFF-state control. Following Cre delivery, robust CD151 depletion was observed, demonstrating efficient activation of both systems (Figure 2A). These data establish that, unlike tRNA-containing constructs (Figure S1A), the mascRNA module does not drive Cre-independent guide transcription, consistent with its lack of canonical internal Pol III promoter elements.

To determine whether the improved editing kinetics reflected enhanced recombination rather than improved sgRNA function, recombined guide cassettes were amplified and loxP recombination efficiency quantified (Figure 2B, Figure S2). Recombination frequencies were comparable between conventional and mascRNA-containing constructs, indicating that enhanced editing performance results from improved sgRNA maturation rather than increased Cre recombination efficiency.

## Discussion

Temporal control of CRISPR perturbation is increasingly important for dissecting dynamic biological processes, modelling transient cellular states, and interrogating essential genes whose constitutive disruption limits interpretation (Zhu et al., 2016; Behan et al., 2019; Norman et al., 2019). Although inducible Cas9 systems provide temporal regulation, they remain constrained by basal activity, variable induction, and the requirement for stable inducible Cas9 cell line generation (Zetsche et al., 2015). Conditional sgRNA activation provides an alternative approach by controlling guide availability while maintaining constitutive Cas9 expression. Cre-loxP-based sgRNA systems are particularly attractive due to their stringent OFF-state control, compatibility with existing Cas9 models, and flexibility in Cre delivery approaches (Jullien et al., 2003; Van den Plas et al., 2003; Chylinski et al., 2019). However, these systems retain a residual 5′ loxP sequence following recombination. Here, we demonstrate that this recombination scar substantially impairs sgRNA performance and that restoration of native guide architecture through mascRNA-mediated processing improves editing kinetics and perturbation penetrance.

The utility of mascRNA as a processing scaffold derives from two key properties: its exploitation of endogenous RNase P and RNase Z processing pathways, which are broadly conserved and active across mammalian cells (Wilusz et al., 2008; Arun et al., 2020), and its lack of intrinsic Pol III promoter activity (Wilusz et al., 2008; Skeparnias et al., 2024). Our observation that tRNA-based processing elements compromise conditional guide fidelity highlights the importance of selecting processing modules that separate RNA maturation from unintended transcriptional activity. mascRNA therefore provides a suitable solution for conditional sgRNA systems requiring both efficient processing and stringent OFF-state control.

The advantages of scarless guide activation extend beyond editing speed. mascRNA-containing constructs achieved more rapid and complete target depletion than conventional loxP-containing guides, reaching levels of perturbation not attained by the scarred sgRNA architecture even at later time points. Delayed or incomplete editing can obscure early molecular responses and reduce sensitivity in time-resolved studies, including investigations of transcriptional adaptation, stress signalling, epigenetic regulation, and synthetic lethality. Importantly, these improvements were achieved without detectable background editing before induction, enabling temporally synchronised perturbation while maintaining OFF-state fidelity.

Guide-to-guide variability remains a major limitation in pooled CRISPR screening applications (Sanson et al., 2018; Behan et al., 2019). Because the impact of the residual 5′ loxP sequence may depend on spacer sequence and local guide architecture, this scar has the potential to introduce additional variability across pooled libraries. By restoring the native sgRNA terminus, mascRNA processing may reduce this source of variability; future genome-wide screens will be required to determine whether this translates into improved guide consistency and more reliable phenotypic scoring.

The modularity of this platform also provides opportunities beyond nuclease-mediated knockout. Scarless conditional guides may be applicable to CRISPR interference, CRISPR activation, base editing, and epigenome engineering systems, where guide abundance and occupancy kinetics influence downstream phenotypes. Furthermore, because activation relies on Cre-mediated recombination, the architecture is compatible in principle with multiple Cre delivery strategies, including transient mRNA electroporation (Van den Plas et al., 2003), recombinant protein delivery (Peitz et al., 2002), and inducible Cre expression systems (Indra et al., 1999).

This flexibility may also prove advantageous for in vivo CRISPR screening applications, where guide activation could be delayed until after cell engraftment or triggered in specific tissues using established Cre driver or inducible Cre systems. Temporally separating cell transplantation from genome editing could minimise engraftment-associated selection effects and enable perturbations to be initiated at defined stages of tumour establishment, disease progression, or tissue homeostasis.

An additional application of this platform is the generation of temporally controlled dual-guide perturbation systems. For example, one guide could be constitutively expressed while a second guide is activated conditionally, enabling interrogation of how the timing of individual gene loss influences the function of a second genetic perturbation. Such designs may be particularly valuable for modelling tumour evolution, genetic interactions, and context-dependent dependencies during tumorigenesis.

Future studies will determine the scalability of this approach in genome-wide pooled screening formats and evaluate its integration with combinatorial perturbation strategies. Temporally synchronised perturbation may provide new opportunities to study adaptive resistance mechanisms, developmental trajectories, and dynamic signalling networks.

In conclusion, we developed a scarless conditional sgRNA platform that combines Cre-loxP-mediated temporal control with endogenous mascRNA processing to eliminate the 5′ loxP-derived sequence that compromises guide activity in existing designs. The system maintains stringent OFF-state control, activates rapidly following Cre induction, and restores editing kinetics comparable to constitutive, unmodified sgRNAs. By providing a modular approach to preserve guide integrity during conditional activation, this platform offers a broadly applicable framework for time-resolved functional genomics, essential gene interrogation, and dynamic CRISPR-based perturbation studies.

## Author contributions

NG and DW conceived the project. NG, CH, LPSD, KS, and AS planned and performed the experiments. CH and CM performed the bioinformatics analysis. NG, CH, DW and LPSD wrote the manuscript. NG, SL, DW and DRT supervised the project. All authors reviewed and approved the manuscript.

## Competing Interest Statement

NG, CH, LPSD, CM and DW are employees of Cancer Research Horizons. SL and DRT are employees of AstraZeneca.

## Acknowledgements

The authors dedicate this work to the memory of Gregory J. Hannon. His impact on the conception of this project and his enduring scientific vision continue to inspire our research. We are very grateful to our colleagues at Functional Genomics Centre for their contributions and useful advice. The authors gratefully acknowledge Cancer Research UK - Cambridge Institute (CRUK-CI) Genomics Facility for their support and assistance in this work.

## Notes

### Summary of Updates

Added Supplementary File 1 with all sgRNA, primer, and other sequences.

