## Supplementary Figures for "Scarless conditional sgRNAs via endogenous mascRNA processing enable rapid and temporally controlled genome editing"

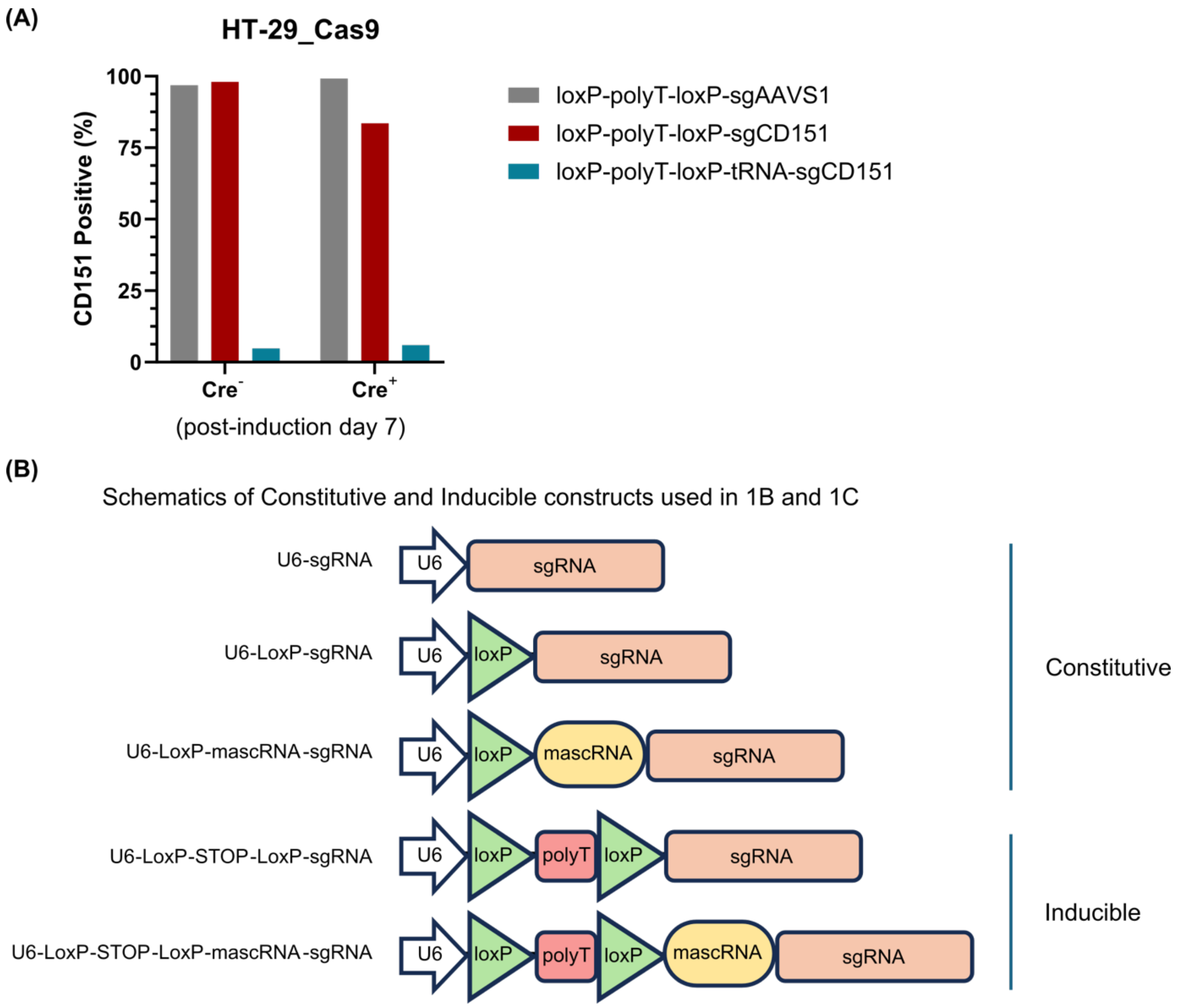


**Figure S1.** (A) CD151 surface expression in HT-29 Cas9 cells transduced with the indicated inducible sgRNA constructs and assessed 7 days after Cre mRNA electroporation or mock electroporation. Canonical tRNA processing modules caused Cre-independent guide activity in the uninduced state. (B) Schematic representation of the constitutive and inducible sgRNA constructs architecture used in this study.

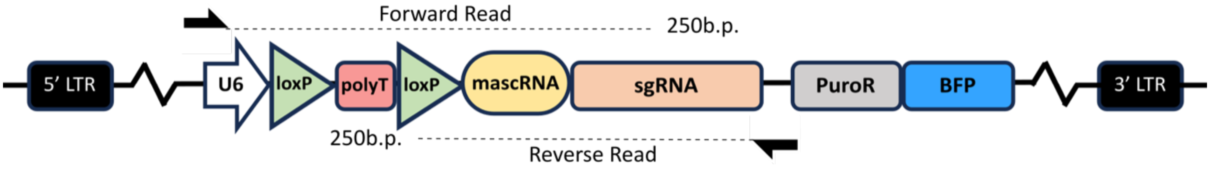


**Figure S2.** Schematic representation of the inducible sgRNA constructs and primer positions used to amplify the integrated guide cassettes for sequencing-based quantification of Cre-loxP recombination.
